# The efficient induction of human retinal ganglion-like cells provides a platform for studying optic neuropathies

**DOI:** 10.1101/2023.07.01.547305

**Authors:** Roxanne Hsiang-Chi Liou, Shih-Wei Chen, Hui-Chen Cheng, Pei-Chun Wu, Yu-Fen Chang, An-Guor Wang, Ming-Ji Fann, Yu-Hui Wong

**Affiliations:** Brain Research Center, National Yang Ming Chiao Tung University, Taipei 112, Taiwan (ROC); Department of Life Sciences and Institute of Genome Sciences, College of Life Sciences, National Yang Ming Chiao Tung University, Taipei 112, Taiwan (ROC); Department of Ophthalmology, Taipei Veterans General Hospital, Taipei 112, Taiwan (ROC); Department of Ophthalmology, School of Medicine, National Yang Ming Chiao Tung University, Taipei 112, Taiwan (ROC); Program in Molecular Medicine, College of Life Sciences, National Yang Ming Chiao Tung University, Taipei, Taiwan (ROC); LumiSTAR Biotechnology, Inc., Taipei 115, Taiwan (ROC)

**Keywords:** Human induced pluripotent stem cells (hiPSCs), induced retinal ganglion cells (iRGCs), direct conversion, ATOH7, BRN3B, SOX4

## Abstract

Retinal ganglion cells (RGCs) are essential for vision perception. In glaucoma and other optic neuropathies, RGCs and their optic axons undergo degenerative change and cell death; this can result in irreversible vision loss. Here we developed a rapid protocol for directly inducing RGC differentiation from human induced pluripotent stem cells (iPSCs) by the overexpression of *ATOH7*, *BRN3B* and *SOX4*. The hiPSC-derived RGC-like cells (iRGCs) show robust expression of various RGC-specific markers by whole transcriptome profiling. A functional assessment was also carried out and this demonstrated that these iRGCs display stimulus-induced neuronal activity, as well as spontaneous neuronal activity. Ethambutol (EMB), an effective first-line anti-tuberculosis agent, is known to cause serious visual impairment and irreversible vision loss due to the RGC degeneration in a significant number of treated patients. Using our iRGCs, EMB was found to induce significant dose-dependent and time-dependent increases in cell death and neurite degeneration. Western blot analysis revealed that the expression levels of p62 and LC3-II were upregulated, and further investigations revealed that EMB caused a blockade of lysosome-autophagosome fusion; this indicates that impairment of autophagic flux is one of the adverse effects of that EMB has on iRGCs. In addition, EMB was found to elevate intracellular reactive oxygen species (ROS) levels increasing apoptotic cell death. This could be partially rescued by the co-treatment with the ROS scavenger NAC. Taken together, our findings suggest that this iRGC model, which achieves both high yield and high purity, is suitable for investigating optic neuropathies, as well as being useful when searching for potential drugs for therapeutic treatment and/or disease prevention.

## INTRODUCTION

Optic neuropathies are a collection of diseases that are characterized by degeneration of retinal ganglion cells (RGCs) and this results in optic nerve damage and visual loss. Common optic neuropathies include glaucoma, ischemic optic neuropathy, toxic optic neuropathy, and so on, which may cause significant visual impairment and economic burden of society [33, 38]. Leber’s hereditary optic neuropathy (LHON) and autosomal dominant optic atrophy (ADOA) are the most frequent forms of hereditary optic nerve disorders. LHON is a mitochondrial genetic disease caused by mutations that affect mitochondrial complex I subunits; this leads to bioenergetic dysfunction and ultimately RGC apoptosis [3]. ADOA causes insidious visual loss due to bilateral degeneration of the optic nerves [22]. Together, these RGC degenerative diseases represent a large proportion of the population of patients suffering from blindness. However, studies of RGC degeneration have been hindered by the lack of feasible and appropriate *in vitro* RGC models. The immortalized RGC-5 cell line has been widely used as a cell culture model to study the neurobiology of RGCs. However, it has recently been re-characterized as having a mouse cone photoreceptor origin. As a result, there is currently no available mammalian RGC cell line for the study of optic neuropathies [19, 20, 34]. Thus it is crucial to establish a robust *in vitro* human RGC model to advance investigations of RGC-related neurodegenerative diseases.

Human induced pluripotent stem cells (hiPSCs) represent a non-ethically disputed and renewable cellular source for RGC generation. In addition, patient-specific iPSCs can be expanded and utilized for RGC differentiation, providing a unique opportunity for the development of *in vitro* RGC models targeting various RGC-related diseases, including hereditary RGC disorders. Recent protocols used to generate hiPSC-derived RGCs have mostly taken advantage of embryoid body (EB) formation and differentiation [15]. These retinal EB cultures resemble the *in vivo* developmental stages of the fetal retinogenesis, and have provided promising methods for obtaining human RGCs for use in the field of vision science [30]. However, due to the fact that retinal organoids generate a heterogenous population of mixed retinal cell types, containing only a limited proportion of RGCs, additional purification procedures are required to isolate the target RGCs. Moreover, organoid culture is time-consuming and cumbersome. Therefore, the establishment of a simplified RGC differentiation protocol using hiPSCs that is highly efficient is important for the advancement of optic neuropathies modeling and for the development of RGC regeneration strategies.

Among various different methods of hiPSC differentiation available, transcription factor (TF)-based cellular conversion has become a commonplace strategy to generate a desired cell type at high efficiency [10]. Such approaches have been developed for a variety of different neuronal subtypes [2, 31, 40, 42, 44]. Here we show that by ectopic expression of three specific and critical TFs that are known to orchestrate RGC fate decision, namely *ATOH7*, *BRN3B* and *SOX4*, hiPSCs can be differentiated into RGC-like cells (iRGCs) at high purity within two weeks. These iRGCs express the typical molecular markers of RGCs, as well as having the appropriate functionalities of RGCs. Moreover, these iRGCs are able to serve as a model for the study of ethambutol (EMB)-induced ocular toxicity. EMB is a first-line medication used in the management of tuberculosis (TB). However, adverse effects, including optic neuropathy, have been reported in approximately 1% of patients taking EMB at the WHO recommended dose level; furthermore, the risk increases substantially if the dose is increased [4, 5]. In this study, we demonstrate that iRGCs undergo dose-dependent cell death after treatment with EMB and that this may be caused by both an increased level of reactive oxidative species (ROS) and an impaired autophagic flux. Our results provided a possible mechanism for EMB-induced ocular toxicity toward RGCs. Furthermore, this human iRGC-based platform promises to be a useful tool that may aid the identification of novel therapeutic insights related to optic neuropathies. Finally, the model system may greatly help future studies that target novel RGC-related treatments and new RGC regeneration strategies.

## MATERIALS and METHODS

### Vector construction

The plasmid pTet-O-Ngn2-puro was a kind gift from Marius Wernig (Addgene plasmid #52047; http://n2t.net/addgene:52047; RRID: Addgene_52047). The *Ngn2* gene cassette was excised using the restriction enzymes EcoRI and XbaI, and the remaining vector served as the backbone for further downstream cloning. Human *ATOH7*, *BRN3B*, *PAX6*, *IRX6*, and *SOX4* were amplified from a reverse transcript of Human Fetal Brain Total RNA (Takara) by high-fidelity PCR using Phusion DNA Polymerases (Invitrogen) with the forward primer carrying EcoRI site and the reverse primer carrying a FLAG tag and XbaI site. These were cloned into the vector pTet-O-T2A-puro. Alternatively, for multiple selection, a zeocin resistance cassette was substituted for the puromycin resistance cassette to obtain the pTet-O-T2A-zeo backbone. For the bicistronic plasmid pTet-O-ATOH7-2A-BRN3B-2A-puro, human *ATOH7* and *BRN3B* were amplified by high-fidelity PCR with primers carrying a myc/FLAG tag and a linking fragment. The two fragments were then linked together using a glycine-serine-glycine (GSG) linker (-GGAAGCGGA-) after a T2A self-cleaving peptide; this is because the GSG linker has been reported to improve the cleavage efficiency of 2A peptides.

Amplicons were cleaned up after PCR using a GenepHlow™ Gel/PCR Kit (Geneaid) and then subjected to restriction enzyme digestion. The digested fragments were gel-purified and then cleaned up using a GenepHlow™ Gel/PCR Kit before ligation into the appropriate vector using T4 ligase (New England Biolabs). The ligated plasmids were transformed into DH5α, next single colonies were selected and these were then validated by PCR. Plasmids were extracted using a Plasmid DNA Midi Kit (Geneaid) and were verified by restriction enzyme mapping and sequencing. The sequences of the primers used are listed in Table S1.

### Lentivirus production

Lentiviruses were introduced by transfection using Lipofactamine 3000 (Invitrogen) into the HEK293FT cells according to the instructions provided by the manufacturer. In total, 5 μg/mL of pCMV-Δ8.91, 1 μg/mL of pCMV-VSV-G and 6 μg/mL of pTetO plasmids were co-transfected into the HEK293FT cells. Supernatants were collected at 24 hrs and 48 hrs after transfection, passed through a 0.45 μm filter (Corning), and then 20-fold concentrated by centrifugation at 4000g/20 min using an Amicon Ultra-15 Centrifugal Filter Unit (3kDa, Millipore). Viral stocks were aliquoted at 200 μL per vial and stored at -80**°**C until use.

### Cell culture

HEK293FT cells were cultured in Dulbecco’s Modified Eagle’s Medium (DMEM-high glucose, Gibco) supplemented with 10% fetal bovine serum (FBS, Biological Industries), MEM non-essential amino acids (NEAA, 1X, Gibco), 1 mM sodium pyruvate (Gibco), penicillin-streptomycin-glutamine (1X, Gibco) and 500 μg/ml Geneticin (G418).

The human induced pluripotent stem cell (hiPSC) line, NTUH-iPSC-02-02 and NTUH-iPSC-01-05 (abbreviated to N2 and N1, respectively), were purchased from Bioresource Collection and Research Center of Food Industry Research and Development Institute, Taiwan. The use of this hiPSC line follows the Policy Instructions of the Ethics of Human Embryo and Embryonic Stem Cell Research guidelines in Taiwan. In addition, approval from the Institutional Review Boards of National Yang-Ming University (Now National Yang Ming Chiao Tung University) was obtained. The cells were maintained on vitronectin (VTN-N, Gibco, USA) coated culture dishes using daily refreshed Essential 8™ medium (E8, Gibco) at 37°C in a 5% CO_2_ incubator. On reaching 90% confluency, the cells were washed once with Dulbecco’s phosphate-buffered saline (DPBS) without calcium and magnesium (Gibco) and then passaged using a splitting ratio of 1:10 in the presence of 0.5 mM EDTA in DPBS.

### Differentiation of iRGC

For RGC differentiation, hiPSCs were infected with lentiviruses encoding ubiquitously driven reverse tetracycline transactivator (rtTA) and Tet-On promoter driven candidate transcription factors sequentially. For lentiviral infection, hiPSCs were seeded onto a 12-well plate the day prior to infection. The medium was then replaced by 800 μL E8 medium with 200 μL concentrated lentiviral stock and 8 μg/mL polybrene on the day of infection and the cells were incubated for 24 hrs. The medium was refreshed every day afterwards and when 90% confluency was reached, the cells were passaged into new culture dishes using 1 μM Y27632 (MedChemExpress). For subsequent differentiation, the infected hiPSCs were seeded at 70% confluency. On the next day, defined as day 0, the differentiation process was started by addition of 2 μg/mL doxycycline. Cells were then cultured in Medium-1: DMEM/F12 (Gibco) supplemented with N-2 (Gibco), 0.1 mM NEAA and 200 ng/mL mouse laminin (Gibco) with various other supplementary substances, namely brain-derived neurotrophic factor (BDNF, 10 μg/mL, PeproTech), ciliary neurotrophic factor (CNTF, 10 μg/mL, PeproTech), basic fibroblast growth factor (bFGF, 5 μg/mL, PeproTech), and forskolin (10 μM, Sigma-Aldrich). On day 1, Medium-1 was refreshed and 1 μg/mL puromycin and 2 μg/mL zeocin were added to allow selection of successfully induced cells. On day 2, the cells were then dissociated using 0.5 mM EDTA in DPBS and replated onto poly-L-lysine- and laminin-coated culture plates for culture in Medium-2: DMEM/F12 + Neurobasal (1:1) supplemented with N-2, B-27, GlutaMAX (Gibco) and 200 ng/mL mouse laminin (Gibco) with supplementary substances. Zeocin selection was continued until day 6.

On day 4 after doxycycline induction, Medium-2 was refreshed with 25 μM AraC to inhibit proliferation of undifferentiated cells. On day 6, after 48 hrs of AraC treatment, Medium-2 was replaced by Medium-3: Neurobasal supplemented with N-2, B27 and GlutaMAX with supplementary substances. Hereafter, the cells were cultured in Medium-3 and medium was half refreshed every 3-4 days until day 15 or day 20, when the cells were used for downstream applications. The supplementary substances included doxycycline (2 μg/mL), brain-derived neurotrophic factor (BDNF, 10 μg/mL, PeproTech), ciliary neurotrophic factor (CNTF, 10 μg/mL, PeproTech), neurotrophin-3 (NT-3, 10 μg/mL, PeproTech), basic fibroblast growth factor (bFGF, 5 μg/mL, PeproTech), insulin (5 μg/mL, PeproTech) and forskolin (10 μM, Sigma-Aldrich); these were added freshly to the medium throughout the whole timespan of the differentiation process. A flow diagram and the culture ingredients used for the iRGC generation are shown in Fig. S2a.

### Immunofluorescence microscopy

The hiPSC-differentiated neurons were cultured on coverslips and subjected to immunofluorescence. Cells were rinsed with PBS and fixed with 4% paraformaldehyde in PBS for 15 min. After fixation, the cells were rinsed with PBS and permeabilized using 0.2% Triton X-100 in PBS for 15 mins. Next, the cells were blocked at room temperature in 3% bovine serum albumin in PBS for 1 hr. Appropriate primary antibodies in the blocking solution were added to the cells, which were incubated at 4°C overnight. After removal of the primary antibodies and rinsing, secondary antibodies in blocking solution together with 1 μg/mL of DAPI were added to cells for 1 hr at room temperature. The antibodies and dilutions used in this study are listed in Table S2. Samples were then mounted on glass slides using mounting solution and examined using a fluorescence microscope (Zeiss).

### Protein extraction and Western blot analysis

Cells were lysed using RIPA lysis buffer (50 mM Tris-HCl pH 7.4, 150 mM NaCl, 1% Igepal-CA630, 0.5% sodium deoxycholate, 0.1% SDS, 2 mM EDTA, 50 mM NaF and protease inhibitor cocktail (Roche). The protein concentrations were determined using a BCA Protein Assay Kit (Pierce). Lysates were denatured at 60°C for 15 min and analyzed by SDS-PAGE and Western blot analysis. For gel electrophoresis, proteins were separated on an 18% SDS-PAGE gel at 100 V for 6 hrs or on 12% SDS-PAGE gel at 100 V for 2 hrs in running buffer (25 mM Tris, 190 mM glycine, 0.1% SDS). Individual gels were electrophoretically transferred to PVDF membranes using 70 V for 2 hrs in transfer buffer (25 mM Tris, 190 mM glycine and 20% methanol). Each membrane was then blocked using 5% skimmed milk and 0.1% Tween 20 in PBS for 1 hr and then probed with primary antibody at 4°C overnight. Membranes were then rinsed with PBST (0.1% Tween 20 in PBS), which was followed by incubation for 1 hr at room temperature with the appropriate secondary antibodies coupled to horseradish peroxidase (HRP). After washing out the secondary antibodies with PBST, the signals were visualized using Immobilon Western Chemiluminescent HRP Substrate (Millipore). GAPDH was used as a loading control.

### RNA extraction, reverse transcription and quantitative real time PCR

Total RNA from hiPSC-differentiated RGC cells was extracted using a Total RNA Mini Kit (Geneaid) according to the user manual. The RNA was reverse transcribed into cDNA using the SuperScript IV reverse transcriptase (Thermo Fisher Scientific) at 55 ℃ for 20 minutes and 80 ℃ for 10 minutes. Transcript levels were quantified by quantitative real-time PCR (qRT-PCR) assays using the appropriate probes from the Universal Probe Library (UPL, Roche). The website service (https://lifescience.roche.com/en_tw/brands/universal-probe-library.html) was used to select the targeted genes, the UPL probes and the primer pairs. Equal amounts of cDNA were mixed with TaqMan™ Master Mix (Roche) and corresponding primers and probes. All qPCRs were conducted at 50 ℃ for 2 min, 95 ℃ for 10 min, and then 40 cycles of 95 ℃ for 15 s and 60 ℃ for 1 min. The threshold crossing value was noted for each transcript and normalized against the corresponding undifferentiated hiPSC control groups and the internal control *RPL13A*. The relative quantification of each mRNA was performed using the comparative CT method. Experiments and data were performed and processed on StepOne Software version 2.3 (Applied Biosystems). The sequences of the primers and probes used are listed in Table S3.

### mRNA library constructing and sequencing

Total RNA of iRGCs from days 7, 14 and 28 were extracted using RNA Mini Kit (Geneaid). mRNA library constructing and sequencing was performed by Azenta Life Sciences company. Briefly, 1 μg of total RNA was used for following library preparation. The poly(A) mRNA isolation was performed using Oligo(dT) beads. The mRNA fragmentation was performed using divalent cations and high temperature. Priming was performed using Random Primers. First strand cDNA and the second-strand cDNA were synthesized. The purified double-stranded cDNA was then treated to repair both ends and add a dA-tailing in one reaction, followed by a T-A ligation to add adaptors to both ends. Size selection of adaptor-ligated DNA was then performed using DNA Clean Beads. Each sample was then amplified by PCR using P5 and P7 primers and the PCR products were validated. Then libraries with different indices were multiplexed and loaded on an Illumina NovaSeq6000 instrument for sequencing using a 2×150 paired-end (PE) configuration according to manufacturer’s instructions.

### Bioinformatics workflow

Sequence read archives (BioProject Number: PRJNA605854, PRJNA413944, PRJNA330714) from published database were downloaded using fastq-dump in SRA toolkit. The fastq format reads underwent quality filter (score >= Q15, qualified percentage > 60%) and adapter trimming using fastp (0.20.1). The remain reads were mapped to Human Genome Reference GRCh38 using STAR (2.7.5a) to generate output in the BAM file format. Expected counts were then estimated using RSEM (1.3.1). Downstream analysis of the expected counts was performed in R Statistical Software (v4.2.1; R Core Team 2022) using DESeq2 (1.36.0). Variance stabilizing transformation, differentially expressed gene (DEG) analysis and principal component analysis were performed using vst, DEseq and plotPCA, respectively, in DESeq2 package. Top 500 up-regulated DEGs, sorted by adjusted p-value, from DESeq2 comparing iRGC on day 7/14/28 to hiPSC were analyzed using Enrichr [39]. Sample distance analysis is performed using the R built-in function - dist. For gene clustering analysis, cell type specific genes were selected from human retina researches [13, 21, 25, 26], and the heatmap was generated using pheatmap(1.0.12).

### Intracellular Ca^2+^ imaging

iRGCs were cultured on coverslips for intracellular calcium imaging. The medium was replaced with loading buffer (150 mM NaCl, 5 mM glucose, 10 mM HEPES, 1 mM MgCl_2_·6H_2_O, 5mM KCl and 2.2 mM CaCl_2_·2H_2_O) containing 2 μM fura-2 or fluo-4. Next the cells were incubated at 37 ℃ for 30 mins, rinsed with loading buffer, and then incubated for a further 20 minutes to allow complete de-esterification of intracellular AM esters. Following this, 1 μM nifedipine and 10 μM ω-conotoxin were added as L-type and N-type voltage-dependent calcium channel antagonists, respectively. In addition, 1 μM tetrodotoxin (TTX) was used as a sodium channel blocker. The coverslips were then loaded on a live-cell imaging culture chamber and the cells were maintained in loading buffer. The chamber was subjected to Ca^2+^ recording at room temperature. Images were acquired through a CCD camera at a time resolution of 1 second per frame. The Ca^2+^ images were recorded before, during, and after a bath application of 70 mM KCl solution at the 50^th^ second of recording. Images were acquired and analyzed using MetaFluor® Fluorescence Ratio Imaging Software. Data are presented as a fluorescence ratio (F340/F380) signal for fura-2, or as ΔF/F0 for fluo-4.

### EMB treatment on iRGCs

EMB of different concentrations ranging from 50 to 2000 μg/ml were added to iRGCs at day 15 after doxycycline induction, and treatment was continued for another three days. Cells were subjected to a viability assay or Western blot analysis on day 20 to 22 after induction. To assess the effects of EMB on autophagy, iRGCs were treated with a saturating concentration of EMB (500 or 1000 μg/ml) with or without 30 or 40 μM of chloroquine (Invitrogen) for 24 hrs. Treated iRGCs were then analyzed by Western blotting using anti-LC3 and anti-p62 antibodies.

### Cell viability assay using alamarBlue^®^

Cultured medium was replaced with fresh medium and a 1/10 volume of alamarBlue^®^ was then added; the cells were incubated at 37°C in a 5% CO_2_ incubator for 4 hrs. Once the color turned pink, the fluorescence was then measured (excitation: 560 nm, emission: 590 nm) using a TECAN Infinite 200. Fluorescence from the wells containing no cells, but treated in the same manner, was measured as the negative control. Cell viability was calculated according to the equation below:

Cell viability (%) = (mean reads of the sample – mean reads of the negative controls) / (mean reads of the sample without EMB treatment – mean reads of the negative controls)

### Monitoring the autophagic process using a Premo™ Autophagy Tandem Sensor RFP-GFP-LC3B Kit

Premo™ Autophagy Tandem Sensor RFP-GFP-LC3B Kits were purchased from Invitrogen. For monitoring the autophagic process in iRGCs treated with EMB, the Baculovirus reagents were added directly into the cultured medium at 24 hrs after EMB treatment and were retained in the medium for 48 hrs to allow transduction. The volume of reagents used was calculated according to the equation below:

Volume of Baculovirus reagents (mL)= (number of cells) × (particles per cells) / 10^8^

After infection with the Baculovirus reagents, the cells were fixed and stained with DAPI. Finally, the fluorescence was visualized using a confocal microscope.

### Flow cytometry to measure intracellular ROS

Human iRGCs were treated with EMB at different concentration ranging from 250 to 1000 μg/mL for 24 hrs or alternatively treated with H_2_O_2_ or Chloroquine for 1 hr. Cells were then dissociated using 0.5mM EDTA into a single cell suspension and incubated in 5 μM of cell-permeant 2’,7’-dichlorodihydrofluorescein diacetate (CM-H_2_DCFDA) dye (Invitrogen) for 30 min at 37°C in the dark. After washing twice with FACS buffer (DPBS with 1% FBS and 0.1% sodium azide), the cells were resuspended in propidium iodide (PI) containing FACS buffer and then analyzed using a Beckman Coulter CytoFLEX analyzer. The results were subjected to further analysis using Flow Jo software (BD Bio sciences). Briefly, the cells were first gated for proper FSC signals, SSC signals and PI-negative signals (for live cells), and then the signals of gated cells were analyzed using a FITC (525/540 nm) filter.

## RESULTS

### Transcription factor-based reprogramming of hiPSC into RGC-like cells

In a landmark study, Zhang et al. demonstrated that expression of neurogenin 2 (NGN2) alone was able to convert hiPSCs into functional induced neurons (iNs) with an almost 100% yield and purity in less than two weeks [44]. By applying a similar reprogramming procedure, we explored the potential of using defined transcription factors (TFs) to bring about the differentiation of RGCs from hiPSCs. In total, seven TFs were initially selected based on their functional roles in the developmental retinogenesis and maintenance of RGCs [6, 8, 16, 23, 27] (Figs. 1a and S1a). The candidate TFs were placed under the control of a Tet operator (TetO) and expressed in hiPSCs via a lentiviral delivery system, along with the constitutive expression of the reverse tetracycline-controlled transactivator (rtTA) (Fig. 1b). The tetracycline-induced expression of the FLAG-tagged or Myc-tagged candidate TFs in the hiPSCs was validated by Western blot analysis (Fig. S1b). Surprisingly, four of the tested TF combinations, *ATOH7*, *ATOH7/BRN3B*, *ATOH7/SOX4*, and *ATOH7/BRN3B/SOX4*, were able to confer typical neuron-like morphologies on hiPSCs resulting in prominent extension of their cell processes (Fig. S2b). By way of contrast, the transduced hiPSCs under other reprogramming conditions failed to survive under the neuronal culturing condition (Fig. S1c). The resulting neuron-like cells also showed expression of the axonal marker SMI312 after two weeks of differentiation (Fig. S2b), which further confirmed the successful neuronal differentiation of hiPSCs by overexpression of these TFs.

**Fig 1.**
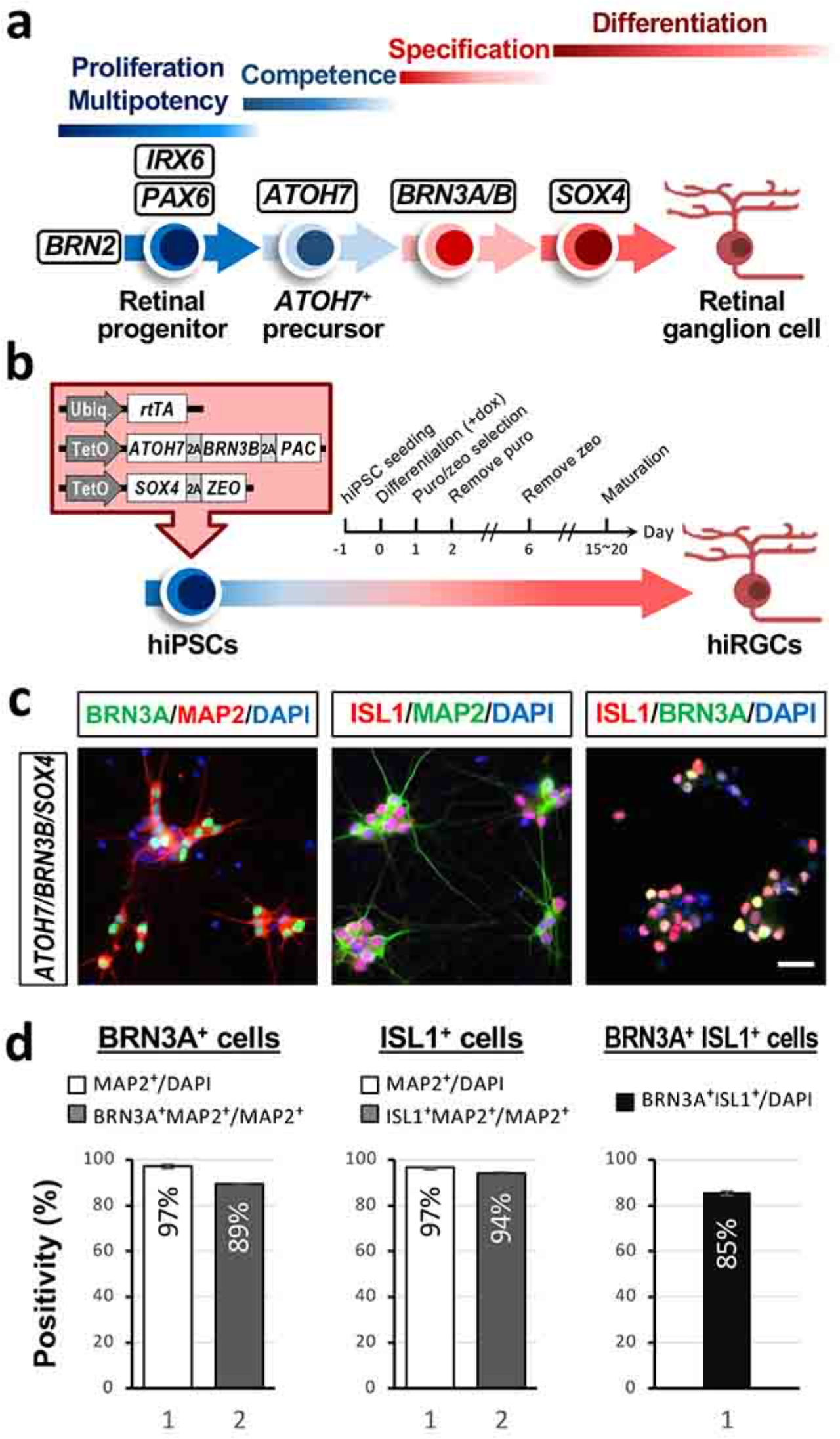
Ectopic expression of transcription factors in hiPSCs induces RGC-like cells. (**a**) A schematic depiction of developmental process used to create the RGCs and the transcription factors that involved. (**b**) Three transcription factors, *ATOH7*, *BRN3B* and *SOX4*, were chosen as reprogramming factors to induce RGCs from hiPSCs. (**c**) Representative images of iRGC cells immunostained for neuronal (MAP2) and RGC (ISL1, BRN3A) markers. Scale bar, 20 μm. (**d**) Quantification of the BRN3A^+^, ISL1^+^, or BRN3A^+^ISL1^+^ cells present among hiPSCs ectopically expressing the candidate genes. Data are presented as means ± SEM (n = 5 batches of independent differentiation).

To further characterize the hiPSC-derived neuron-like cells formed under the four aforementioned reprogramming conditions, we adapted the culturing conditions for rodent primary RGCs according to previous studies in order to further facilitate the maturation and survival of our putative human iRGCs [37]. One day after induction, the transduced hiPSCs underwent positive selection using puromycin and zeocin for two days, and then were replated onto a coating of poly-L-lysine and laminin; this was followed by culture in neuronal medium in the presence of brain-derived neurotrophic factor (BDNF), ciliary neurotrophic factor (CNTF), basic fibroblast growth factor (bFGF) and forskolin supplementation (Fig. S3a). The neuron-like cells produced under these conditions were subjected to immunocytochemistry after two weeks of differentiation in order to examine the expression of the important RGC markers BRN3A and ISL1. While all four reprogramming conditions gave rise to neuron-like cells at a similar efficiency (MAP2^+^/DAPI^+^ cells >90%), the *ATOH7*-expressing and *ATOH7/SOX4*-expressing hiPSCs generated proportionately fewer BRN3A-positive (71.4 ± 0.8% and 75.5 ± 0.6%, respectively) and ISL1-positive neurons (64.7 ± 2.4% and 67.2 ± 1.4%, respectively), with most of these cells appearing unhealthy. On the other hand, the combined overexpression of [*ATOH7* and *BRN3B*] or [*ATOH7*, *BRN3B* and *SOX4*] in hiPSCs resulted in the highest efficiency when generating iRGCs, with the majority of the MAP2-positive cells being BRN3A-positive (87.6 ± 0.3% and 89.5 ± 0.1%, respectively) or ISL1-positive (97.2 ± 0.3% and 94.1 ± 0.4%, respectively) (Figs. S3b, c and d). Furthermore, *ATOH7/BRN3B* and *ATOH7/BRN3B/SOX4* gave rise to overlapping BRN3A and ISL1 expression in the majority of the population (87.2 ± 2.0% and 85.3 ± 0.9%, respectively), which further confirmed the identity of these cells as RGC-like cells (Fig. S3e). Interestingly, although exhibiting comparable efficacy in terms of RGC differentiation, the *ATOH7/BRN3B*-expressing iRGCs appeared to be in a poor condition that the cells form large compact clusters with noticeable cell death in the inner layer. Collectively, the TF combination of *ATOH7*, *BRN3B* and *SOX4* seemed to have the highest potential for generating high purity RGC-like cells from hiPSCs. These RGC-like cells induced by the three TFs were hereafter termed iRGC or iRGC-*ATOH7/BRN3B/SOX4*.

### Characterization of the hiPSC-derived RGCs (iRGCs)

To further characterize the identity of these iRGCs, RT-qPCR analysis was carried out after two weeks of differentiation in order to measure the mRNA expression levels of two neuronal markers, namely *RBFOX3* and *TUBB3*, and various canonical RGC markers, including *ATOH7*, *BRN3A*, *BRN3B*, *EBF1*, *ISL1*, *RBPMS*, *SOX4*, and *SOX11*. Endogenous levels of *ATOH7*, *BRN3B* and *SOX4* were detected using primers targeting the 5’ or 3’ untranslated region of these genes. Both iRGC-*ATOH7/BRN3B* and iRGC-*ATOH7/BRN3B/SOX4* showed elevated expression levels of *RBFOX3*, *TUBB3*, and all of the RGC markers except for *RBPMS* when they were compared to the expression levels in hiPSCs; however, only iRGC-*ATOH7/BRN3B/SOX4* exhibited a significant increase in expression of *RBPMS* (Fig. 2a). In addition, since the majority of RGCs are glutamatergic neurons, the expression of vesicular glutamate transporters 1 and 2 (*SLC17A7* and *SLC17A6*, respectively) were also examined. The expression levels of *SLC17A7* and *SLC17A6* were significantly higher in both iRGC-*ATOH7/BRN3B* and iRGC-*ATOH7/BRN3B/SOX4* as expected, while the expression level of the stem cell marker *OCT4* was, as expected, lower (Fig. 2a). In conclusion, the forced expression of *ATOH7*, *BRN3B*, and *SOX4,* together with the addition of appropriate growth factors, can efficiently convert hiPSCs into a RGC lineage within two weeks.

**Fig 2.**
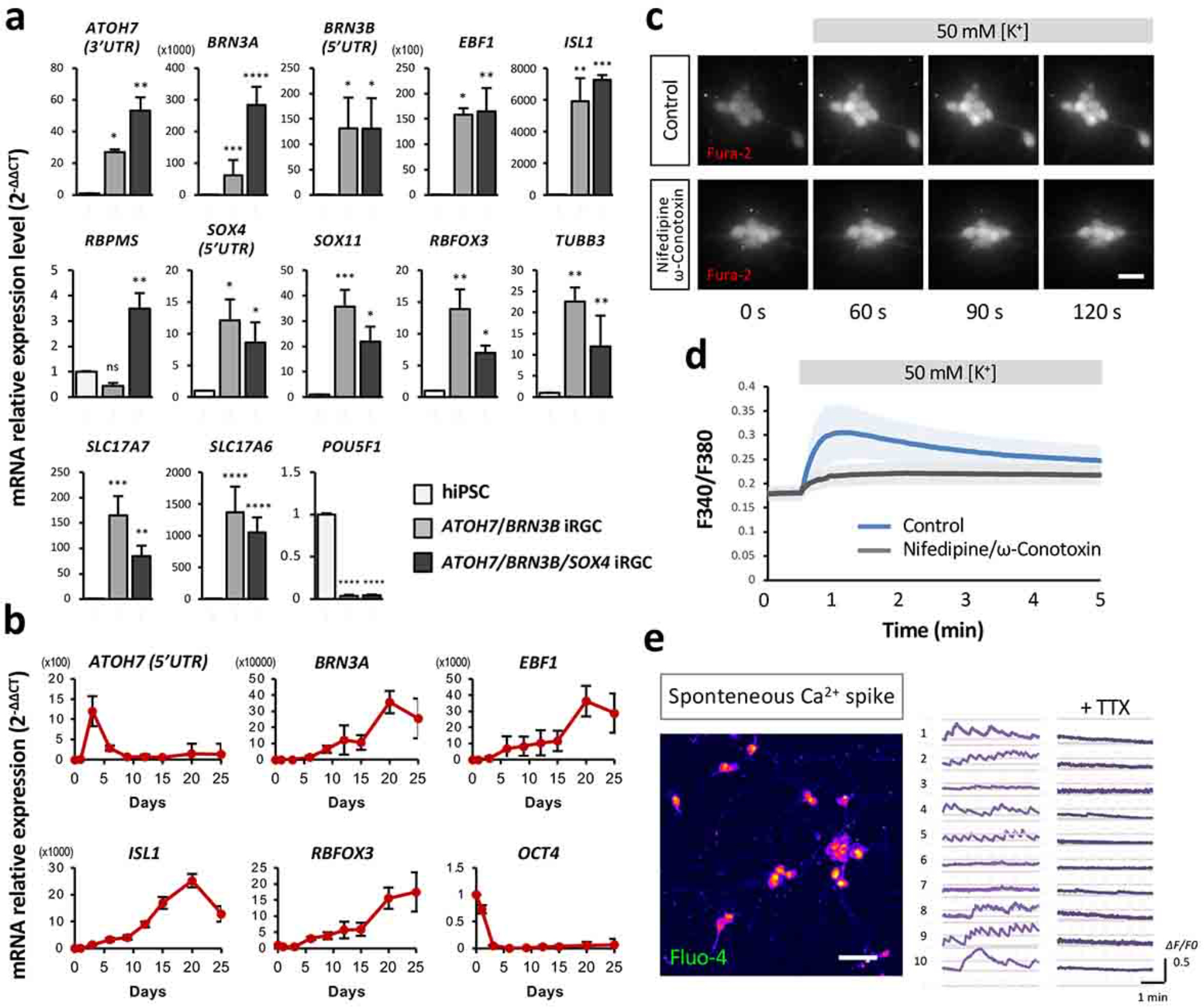
iRGC-*ATOH7*/*BRN3B/SOX4* expresses consensus RGC markers and shows neuronal activity. (**a**) Expression levels of the general RGC markers (*ATOH7*, *BRN3A*, *BRN3B*, *EBF1*, *ISL1*, *RBPMS*, *SOX4* and *SOX11*), the stem cell marker (*OCT4*), the neuronal markers (*RBFOX3* and *TUBB3*), and the glutamatergic neuron markers (*SLC17A7* and *SLC17A6*) in iRGCs at day 15 of induction measured by RT-qPCR. Fold change was calculated using the ΔΔCT method with the corresponding undifferentiated hiPSC groups and *RPL13A* as the controls. The endogenous expression of *ATOH7*, *BRN3B*, and *SOX4* was quantified with primer pairs located in the untranslated region (5’UTR or 3’UTR). Data are presented as means ± SEM. Statistical analysis was carried out using one-way ANOVA (n = 5 batches of independent differentiation; *: *p* < 0.05, **: *p* < 0.01, ***: *p* < 0.001). (**b**) Time course of mRNA expression levels of the key RGC markers (*ATOH7, BRN3A, EBF1, ISL1*) and the stem cell marker (*POU5F1*) by RT-qPCR (normalized against a reference gene *RPL13A)*. Data are presented as the means of four batches of iRGC induction together with the SEM. (**c**) Representative images showing an example of [Ca^2+^]i transients following application of high-K^+^ solution (50 mM) to iRGCs at D15 loaded with the Ca^2+^ indicator Fura-2-AM. Scale bar, 20 μm. (**d**) The time-lapse changes in fluorescence intensity produced by high-K^+^ application. Traces show the mean intensity (F340/F380) of each time point in the presence (black) or absence (blue) of inhibitors. Data are presented as means ± SEM (n = 18-20 cells). (**e**) A representative image showing an example of spontaneous [Ca^2+^]i transients in iRGCs loaded with the Ca^2+^ indicator Fluo-4-AM at D21. Scale bar, 50 μm. The time-lapse changes in fluorescence intensity of each iRGC were recorded at 1 Hz before and after adding 1 μM TTX, and shown as ΔF/F0 (right).

We next assessed the dynamics of RGC marker expression throughout the differentiation process of iRGC-*ATOH7/BRN3B/SOX4* by RT-qPCR analysis. The expression levels of *ATOH7*, *BRN3A*, *EBF1*, *ISL1*, *RBFOX3* and *OCT4* were examined on days 0, 1, 3, 6, 9, 12, 15, 20 and 25 of differentiation. Endogenous *ATOH7* expression was found to be highest after the onset of the induction, reaching its peak on day 3 and then declining; this corresponds with the fact that *ATOH7* is expressed in RGC precursor cells and is known to orchestrate the initiation of the RGC differentiation program [1]. Similarly, the RGC markers *BRN3A*, *EBF1*, and *ISL1* started to be expressed after iRGC induction and reached their maximum levels on day 20. The neuronal marker *RBFOX3* was found to be elevated throughout the differentiation process and continued to be steadily expressed after day 20. On the other hand, the stem cell marker *OCT4* was barely detectable on day 3 and thereafter (Fig. 2b). Collectively, our iRGC differentiation protocol demonstrated that there was rapid differentiation of the hiPSC-derived RGCs, with the majority of the iRGCs reaching maturation after day 15 of differentiation.

### Functional characterization of the iRGCs

After evaluating the molecular identity of iRGC-*ATOH7/BRN3B/SOX4*, we next examined whether these iRGCs showed any general neuronal functions. An increase in intracellular Ca^2+^ influx in response to depolarization is one of the distinctive features of functional neurons, and therefore we used a ratiometric fluorescent calcium indicator, Fura-2-AM, to monitor the Ca^2+^ response of iRGC-*ATOH7/BRN3B/SOX4* upon depolarization. The day 15 iRGCs were subjected to treatment with a 50 mM potassium solution to depolarize the neurons and the intensity of Fura-2 fluorescence induced by 340 nm and 380 nm excitation was recorded; these results are presented as a F340/F380 ratio. Upon depolarization, there was an increase in both intracellular F340 and the F340/F380 ratio, suggesting that there was an increase in intracellular Ca^2+^ in response to depolarization (Figs. 2c and d). To further explore the functionality of the Ca^2+^ response of the iRGCs, nifedipine and ω-conotoxin were applied to the cells as L-type and N-type calcium channel blockers, respectively. The voltage-gated calcium influx was abolished in iRGC-*ATOH7/BRN3B/SOX4* after the administration of these calcium channel blockers, and this was manifested by a failing to increase either intracellular F340 or the F340/F380 ratio. To further examine the neuronal maturity and functionality of the iRGCs, spontaneous neuronal activity was evaluated after 3 weeks of differentiation using another calcium indicator, Fluo-4-AM. Spontaneous firing and the transmission of the action potentials were observed in the iRGCs, as visualized by the cadenced rise-and-fall of Fluo-4 fluorescence intensity over the observation period (Fig. 2e). Such Ca^2+^ events were abolished by the addition of tetrodotoxin (TTX), a sodium channel blocker that inhibits the firing of action potentials in neurons (Fig. 2e, right panel). With the presence of robust activity in terms of both induced and spontaneous neuronal responses, we have confirmed that our iRGCs at the 3-week-differentiation time point have the capability of establishing an integrated neuronal network that possess the electrophysiological characteristics of and the chemical functionality of a RGC model system. Altogether, these results show that the iRGC-*ATOH7/BRN3B/SOX4* has neuronal features comparable to *in vivo* RGCs; these include mature calcium channels and spontaneous neuronal activity; furthermore, these are present after only a short differentiation process of two to three weeks.

### Transcriptome profiling confirms iRGC identity

Next, we performed time-course RNA-seq experiments to characterize iRGC identity transitions during *ATOH7/BRN3B/SOX4*-based differentiation. The results from principal component analysis (PCA) showed distinct clustering of samples, derived from three independent hiPSC lines (N1, N2 and Allele), across different time points of differentiation (week 0, 1, 2 and 4) (Fig. 3a). The gene ontology (GO) analysis revealed that the genes highly expressed in iRGCs on week 1 and 2 of differentiation (iRGC_W1/W2) were enriched for neural development, such as nervous system development and axonogenesis, while those strongly expressed in iRGC_W4 were predominantly associated with synaptogenesis and neuronal functions such as synaptic vesicle endo- and exocytosis and neurotransmitter secretion (Fig. 3b). Further gene enrichment analysis using ENRICHR and the tissue expression library confirmed a stepwise differentiation resulting in the “retina” as the top category by weeks 1, 2, and 4 [39]. The gene expression heatmap analysis also demonstrated consistent cell type and stage-specific gene expression with the progression of hiPSC-to-RGC differentiation (Figs. 3d and S5a). Accordingly, pluripotency-associated genes *POU5F1(OCT4)* and *NANOG* were downregulated by day 7, and by weeks 2 and 4, cells expressed the canonical RGC markers including *DCX, GAP43, NEF1, NEFM, POU4F1, SNCG, STMN2, THY1* and *TUBB3*. However, it was noted that throughout the entire differentiation process, there was no significant induction of genes involved in eye-field specification or retinal progenitor generation, as the direct conversion from iPSCs to iRGCs was based on the expression of three lineage-specific TFs and might have skipped the intermediate steps that occur during early RGC development. Additionally, markers of other main cell types in the retina, such as amacrine cells, bipolar cells, rod and cone photoreceptors, Müller glia, and pigment epithelium, were not present throughout the differentiation process. These findings demonstrate that iPSCs can be directly differentiated into iRGCs without passing through the retinal progenitor phase.

**Fig 3.**
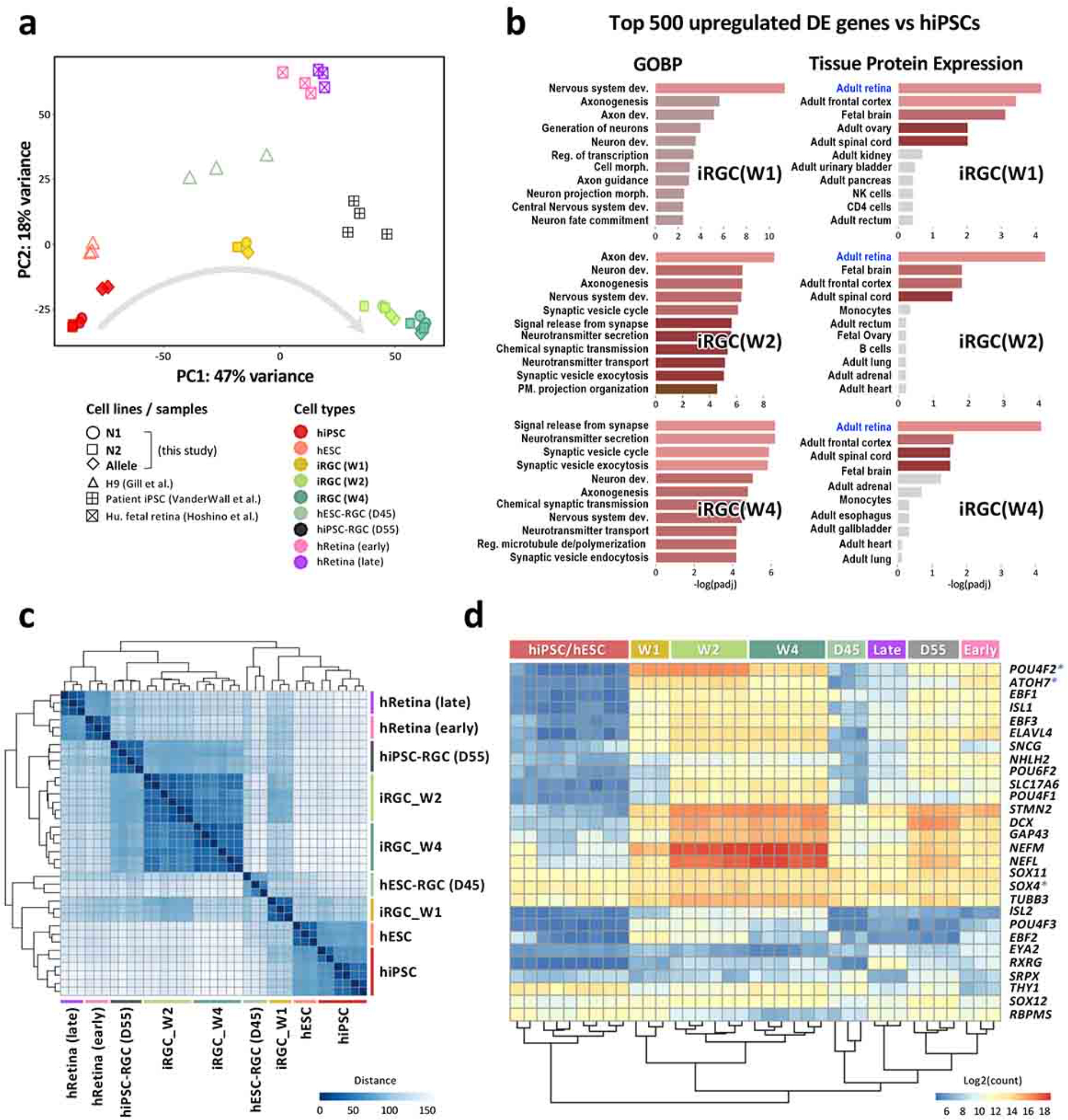
Transcriptomic analysis of iRGCs during the differentiation process. (**a**) Principal component analysis (PCA) of all samples by whole transcriptome sequencing of protein coding genes. The profiles of iRGCs at week 1, 2 and 4 derived from three hiPSC lines with different genetic backgrounds (N1, N2 and Allele) were merged with the dataset of RGCs generated through alternative methods (hiPSC/ESC-RGC) and those of human retina tissues during weeks 8 and 20 of gestation (hRetina_early and hRetina_late, respectively). Each spot represents one independently differentiated cell batch, and each cell type is coded with different colors and shapes. (**b**) Gene enrichment analysis was performed on the top 500 differentially upregulated genes, sorted by adjusted p-value, in iRGCs versus hiPSCs (W0). The eleven most significant groups are presented in terms of gene ontology biological process (GOBP, left panel) and tissue protein expression (right panel). The analysis reveals “retina” as the top hit on week 1, 2, and 4 of differentiation. Gray bars indicate no significant expression (padj > 0.05). (**c**) A heatmap showing the Euclidean distances between samples clustered using unbiased ordering of samples based on sample similarity. (**d**) A heatmap depicting the relative expression of RGC marker genes in all samples. The Tet-on inducible genes are marked with asterisk (*).

To further validate the phenotype of our iRGCs, we compared the transcriptome profile with those generated by alternative methods (hiPSC/ESC-RGC) and those previously reported for human retina tissues during weeks 8 and 20 of gestation (hRetina_early and hRetina_late, respectively) [11, 12, 35] (Fig. S4). The PCA was performed to assess the similarity among the cell types and showed clear clustering of samples (Fig. 3a). Based on the first principal component (PC1), the samples seemed to demonstrate cellular maturation progressing from left to right; on the left is the pluripotent stem cell, and on the right is closer to the mature RGC type. Euclidean distance analysis, which provided an unbiased ordering of the samples, also showed the similarity between our iRGCs and those obtained through EB-based differentiation (Fig. 3c). Furthermore, a clustered heatmap of selected RGC marker genes revealed that our iRGC cells at week 2 and week 4 had a similar expression pattern to the human fetal retina and hiPSC-RGCs derived from EB-mediated differentiation at day 55 (Fig. 3d). Importantly, differential analyses between our iRGC cells on week 4 and hESC-RGCs at day 45 revealed that the most significantly upregulated DEGs in iRGC_W4 were related to the GO terms of neurotransmitter secretion and synaptic transmission (Fig. S5b). These transcriptome profiling results thus confirm that the directly-converted iRGCs on week 4, which expressed a panel of markers associated with RGCs and neuronal function, resemble EB-mediated differentiated RGCs at week 8 (day 55) and the human fetal retina.

### iRGC-*ATOH7/BRN3B/SOX4* as a human RGC model for drug-induced optic neuropathy

After confirming the RGC identity and functionality of iRGC-*ATOH7/BRN3B/SOX4*, we next tested the potential of these iRGCs as a model for investigating RGC-related diseases. EMB-induced optic neuropathy is a well-known complication that results from the use of an anti-tuberculosis drug. Ethambutol is considered the least toxic drug available when treating tuberculosis; however, the occurrence of optic neuropathy as a side effect of EMB treatment has been identified, and this puts five million patients at the risk of irreversible blindness each year [18, 28]. The exact mechanism by which the ocular neurotoxic effect of EMB occurs is yet to be identified. Therefore we exploited iRGC-*ATOH7/BRN3B/SOX4* as a human RGC model to investigate the possible mechanisms behind EMB ocular neurotoxicity.

To investigate the neurotoxic effect of EMB on human RGC cells, different concentrations of EMB were used to treat iRGC-*ATOH7/BRN3B/SOX4* for 72 hrs on day 20 (D20) of differentiation. EMB could be seen to cause significant cell death in a dose-dependent manner as assessed by the alamarBlue cell viability assay (Fig. 4a); this represents a clinically relevant dose-related RGC death similar to that which occurs in EMB-treated patients [9]. The expression level of the apoptosis protein, cleaved caspase-3, was found to significantly increased in a concentration-dependent manner on EMB treatment for 24 hr, while total caspase-3 remained unchanged (Figs. 4c and d). These results indicated that the caspase-3-associated apoptosis pathway was activated following EMB treatment in our iRGC-*ATOH7/BRN3B/*SOX4 model system.

**Fig 4.**
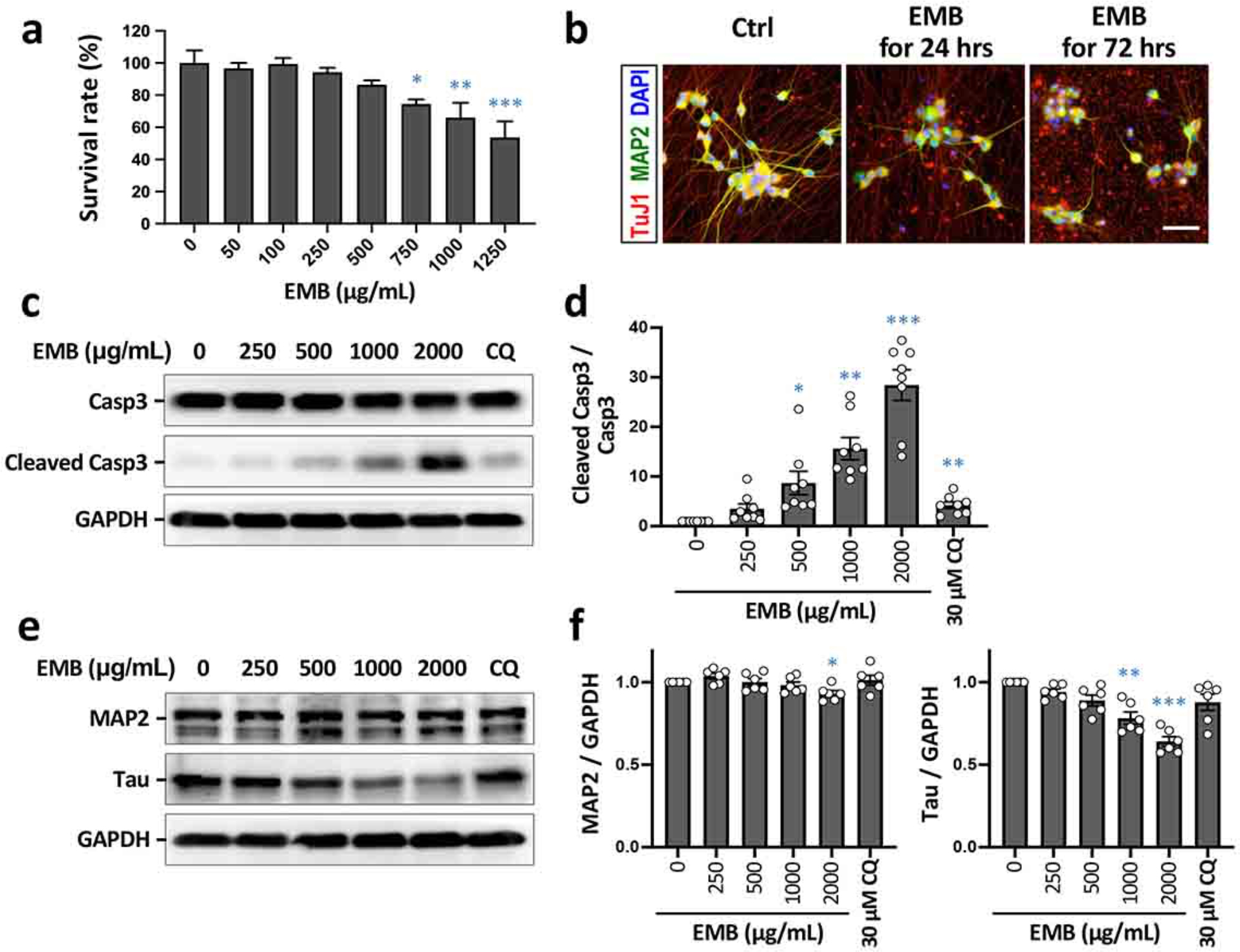
EMB induces neuronal death in iRGC-*ATOH7*/*BRN3B/SOX4*. (**a**) Cell viability assays using alamarBlue® after treatment of iRGCs with various concentrations of EMB for 72 hrs at D20. Values are presented as mean ± SEMs from 4 independent experiments (*: *p* < 0.05, **: *p* < 0.01, ***: *p* < 0.001 versus control by one-way ANOVA and Dunnett’s multiple comparisons test). (**b**) Cell morphological change of the iRGCs after EMB treatment for 24 or 72 hrs and after immunolabeling for MAP2 and TuJ1. Scale bar, 50 μm. (**c**) Expression levels of the apoptotic marker, caspase-3, in iRGCs at D21 using Western blot analysis. Glyceraldehyde 3-phosphate dehydrogenase (GAPDH) was used as a protein loading control. (**d**) Quantification of the protein levels in (**c**). The intensity of each signal was normalized against the control (EMB = 0). Data represent mean ± SEM (n = 8 independent batches of differentiation; *: *p* < 0.05, **: *p* < 0.01, ***: *p* < 0.001 by one-way ANOVA and Dunnett’s test). (**e**) Protein expression levels of MAP2 and Tau in iRGCs at D21 using Western blot analysis. (**f**) Quantitative results of (**e**). The intensity of each signal was normalized against the control (EMB = 0). Data represent mean ± SEM (n = 6 independent batches of differentiation; *: *p* < 0.05, **: *p* < 0.01, ***: *p* < 0.001 by one-way ANOVA and Dunnett’s test).

Following EMB treatment, morphological changes occurred in the iRGCs, including cell shrinkage, increased cell detachment, and a neurodegenerative phenotype involving neurite fragmentation; the changes were compared to untreated control cells and were observed using bright-field microscopy. To ascertain whether EMB is able to cause neurite degeneration in the iRGC-*ATOH7/BRN3B/SOX4* model system, we performed immunostaining for a neuronal marker TuJ1 (neuron-specific Class III β-tubulin). As shown in Fig. 4b, the degenerated and fragmented neurites showed a TuJ1 positive signal with noticeable blebbing after 1000 μg/mL EMB treatment for 24 hr. Intact neurite outgrowth were barely visible following EMB treatment at the same dosage for 72 hr. In the Western blot results, the protein level of Tau, a neuronal microtubule-associated protein enriched in axons, was significantly decreased in iRGC-*ATOH7/BRN3B/SOX4* cells in a dose-dependent manner (Figs. 4e and f). Apart from the axons, EMB also caused significant damage to the dendrites, with immunochemical evaluation of the somatodendritic marker MAP2 being used to show a significant reduction in this marker (Fig. 4b), although this was not statistically significant in the Western blot results (Fig. 4e). Altogether, these results indicated that EMB induces significant neurite degeneration and apoptosis in human RGCs and that this occurs in a dose-dependent and time-dependent manner.

### EMB impairs autophagic flux in iRGCs by blocking autophagosome-lysosome fusion

To further understand how EMB causes such adverse effects on iRGCs, we next endeavored to unearth the mechanism(s) behind EMB neurotoxicity. In previous studies, EMB has been shown to impair autophagic flux and to cause apoptosis in the both the human Huh-7 cell line and in rodent retinal neurons [14, 41]; this suggests that dysregulation of autophagy in RGCs might be one potential cause of EMB-induced optic neuropathy. To further investigate this possibility, we treated D20 iRGC-*ATOH7/BRN3B/SOX4* with non-lethal concentrations of EMB for 24 hrs, then evaluated the protein levels of two autophagy markers, LC3 and p62, by Western blot analysis. Both lipidated LC3-II and lipidated p62 protein were significantly elevated in iRGCs treated with higher doses of EMB (1000 μg/mL and above for LC3-II; 500 μg/mL and above for p62, Figs. 5a, b and c). Both LC3-II and p62 are selectively degraded during autophagy, therefore, an increased accumulation of both of these proteins points towards an impairment in autophagic degradation [43]. To further investigate this increase in LC3-II and p62 proteins in the iRGCs and to determine whether it is due to EMB-induced autophagic impairment, rather than autophagic activation, an autophagy inhibitor, chloroquine (CQ), was added along with the EMB for 24 hrs. The addition of CQ did not further increase the levels of LC3-II and p62 in EMB-treated cells (Figs. 5d, e and f), which fails to support the possibility that EMB increases autophagic activation. Taking these results together, it would seem that EMB at high concentrations results in insufficient autophagic degradation, which then leads to the dose-dependent cell death of the treated iRGCs.

**Fig 5.**
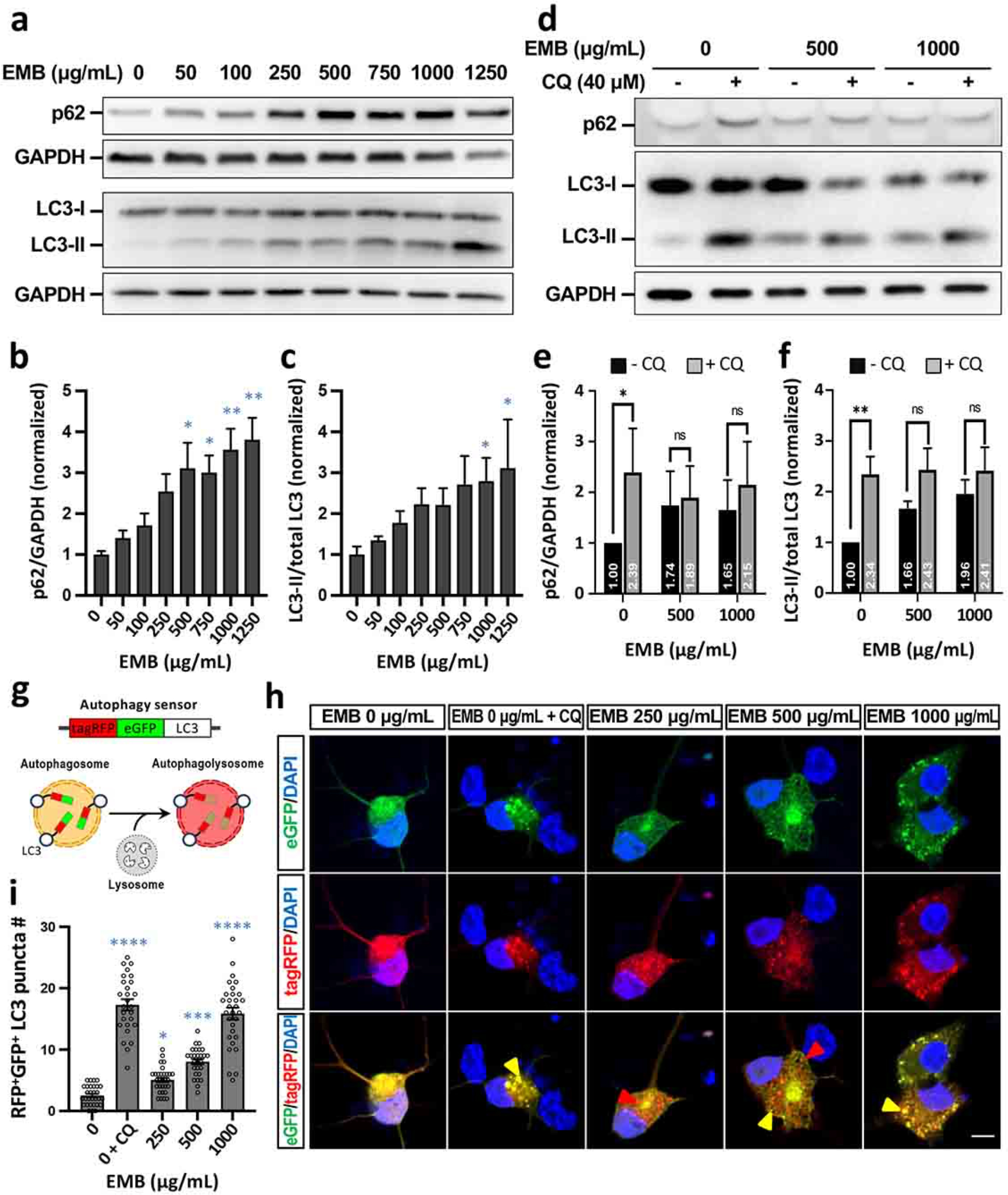
EMB blocks autophagic flux in iRGC-*ATOH7*/*BRN3B/SOX4*. (**a**) The conversion of LC3-I to LC3-II and p62 accumulation after EMB treatment for 24 hrs were analyzed using Western blotting. Blots are representative of the three independent experiments and GAPDH was used as the sample loading control. (**b** and **c**) Quantitative analysis of p62 (**b**) and LC3-II (**c**) levels relative to GAPDH and total LC3, respectively, was performed and the values were normalized against the control group. Data represent mean ± SEM of readings from 3 independent experiments (*: *p* < 0.05, **: *p* < 0.01 versus control (EMB = 0) by one-way ANOVA and Dunnett’s test). (**d**) EMB-induced LC3 and p62 level changes in the presence or absence of CQ were analyzed using Western blotting. (**e** and **f**) Quantitative results of (**d**). Data represent mean ± SEM of readings from three independent experiments (*: *p* < 0.05, **: *p* < 0.01, ns: not significant by two-way ANOVA). (**g**) A schematic depiction of the mechanism of tagRFP-eGFP-LC3. (**h**) iRGCs at D20 were infected with RFP-GFP-LC3 baculovirus. One day post infection the cells were treated with various concentrations of EMB or 40 μM CQ for 24 hrs, fixed and imaged by fluorescence microscopy. The yellow arrowheads indicate an RFP^+^GFP^+^ autophagosome, while the red arrowheads indicate an RFP^+^ autophagolysosome. Scale bar, 10 μm. (**i**) Quantitative results of the number of RFP^+^GFP^+^ LC3 puncta in (**h**). Data represent mean ± SEM from 27-30 iRGCs in each group of three independent experiments (*: *p* < 0.05, ***: *p* < 0.001, ****: *p* < 0.0001 versus control (EMB = 0) by one-way ANOVA and Dunnett’s test).

Accumulation of both LC3-II and p62 implies a deficiency in the lysosome-dependent degradation, which can either be a result of blockage of lysosome-autophagosome fusion or of the inhibition of lysosomal proteases. To elucidate whether EMB interferes with the fusion between autophagosome and lysosome, a tagRFP-eGFP-LC3 tandem fluorescent probe was used to monitor autophagic flux in the EMB-treated iRGCs (Fig. 5g). Upon lysosome-autophagosome fusion, the fluorescence of the eGFP probe is quenched due to an acidic environment whereas that of tagRFP is stably present. In such circumstances, the autophagosome puncta will become eGFP^-^/tagRFP^+^ (red) upon fusion with lysosomes, while the ones that fail to fuse will remain eGFP^+^/tagRFP^+^ (yellow) [17]. The results for the D20 iRGC-*ATOH7/BRN3B/SOX4* cells showed an increased number of red LC3 puncta when treated with 250 μg/mL of EMB for 24 hr (Fig. 5h, red arrowheads), which indicates that there is an ongoing autophagic flux and intact lysosome-autophagosome fusion. However, as the EMB treatment was scaled up to 500 μg/mL, yellow LC3 puncta began to manifest, showing the autophagosome puncta were failing to undergo fusion with lysosomes. As comparable to the CQ-treated control group, treatment with 1000 μg/mL of EMB resulted in a massive increase in yellow LC3 puncta; almost no red puncta were able to be observed under this condition (Fig. 5h, yellow arrowheads). In conclusion, EMB at a high concentration interferes with fusion between lysosomes and autophagosomes in iRGCs, as indicated by the increased ratio of eGFP^+^/tagRFP^+^ LC3 puncta (Fig. 5i). The EMB-induced impediment to lysosome-autophagosome fusion thereafter leads to insufficient autophagic degradation, which in turn results in adverse effects on the iRGCs such as neurite degeneration and significant cell death.

### EMB-induced cell death involves ROS generation in iRGCs

In addition to autophagy dysregulation, intracellular reactive oxygen species (ROS) production by mitochondria has also been reported to be closely related to apoptosis in various cell types [24, 29]. Therefore, we investigated whether intracellular ROS activation was involved in EMB-induced iRGC death. After iRGCs (D20) were treated with EMB for 24 hr, ROS was measured using DCF-DA dyes and flow cytometry (Fig. 6a). We found that EMB treatment significantly enhanced intracellular ROS levels in a dose-dependent manner compared to the untreated controls (Fig. 6b). Hydrogen peroxide (500 μM H_2_O_2_) was used as a positive control to increase ROS levels in the cells, while the CQ (30 mM), as a negative control, failed to increase ROS. These findings indicate that the autophagy defect caused by CQ has no effect on ROS production. Furthermore, EMB-induced autophagy impairment and the increased ROS levels may not necessarily be linked. As expected, the ROS scavenger NAC (antioxidant N-acetyl-L-cysteine) markedly blocked EMB-induced ROS generation in iRGCs (Fig. S6b). When assessed using the alamarBlue assay, NAC co-treatment showed a significant recovery of EMB-induced cell death (Fig. 6c). Notably, NAC treatment (500 μM) alone did not alter ROS levels and had no toxic effect on iRGC viability (Fig. S6). To further confirm the beneficial effect of the antioxidant on EMB-induced cell death, we examined the apoptosis marker, cleaved caspase-3, in response to the treatment with EMB, NAC, or the two combined. As shown in Fig. 6e, EMB treatment for 24 hr increased the level of cleaved caspase-3 in iRGCs, while NAC treatment, remarkably, reduced the level of cleaved caspase-3 when iRGCs were treated with 250-1000 μg/mL of EMB. In addition, NAC significantly mitigated neurite degeneration caused by EMB, as evidenced by increases in MAP2 and TuJ1 staining (Fig. 6d). Western blot analysis also revealed that the expression level of Tau protein was restored under NAC co-treatment (Figs. 6g and h). Collectively, these results suggested that EMB-induced ROS generation causes neurite degeneration and apoptosis of human RGC cells, and that antioxidants such as NAC are capable of alleviating these adverse effects to some degree.

**Fig 6.**
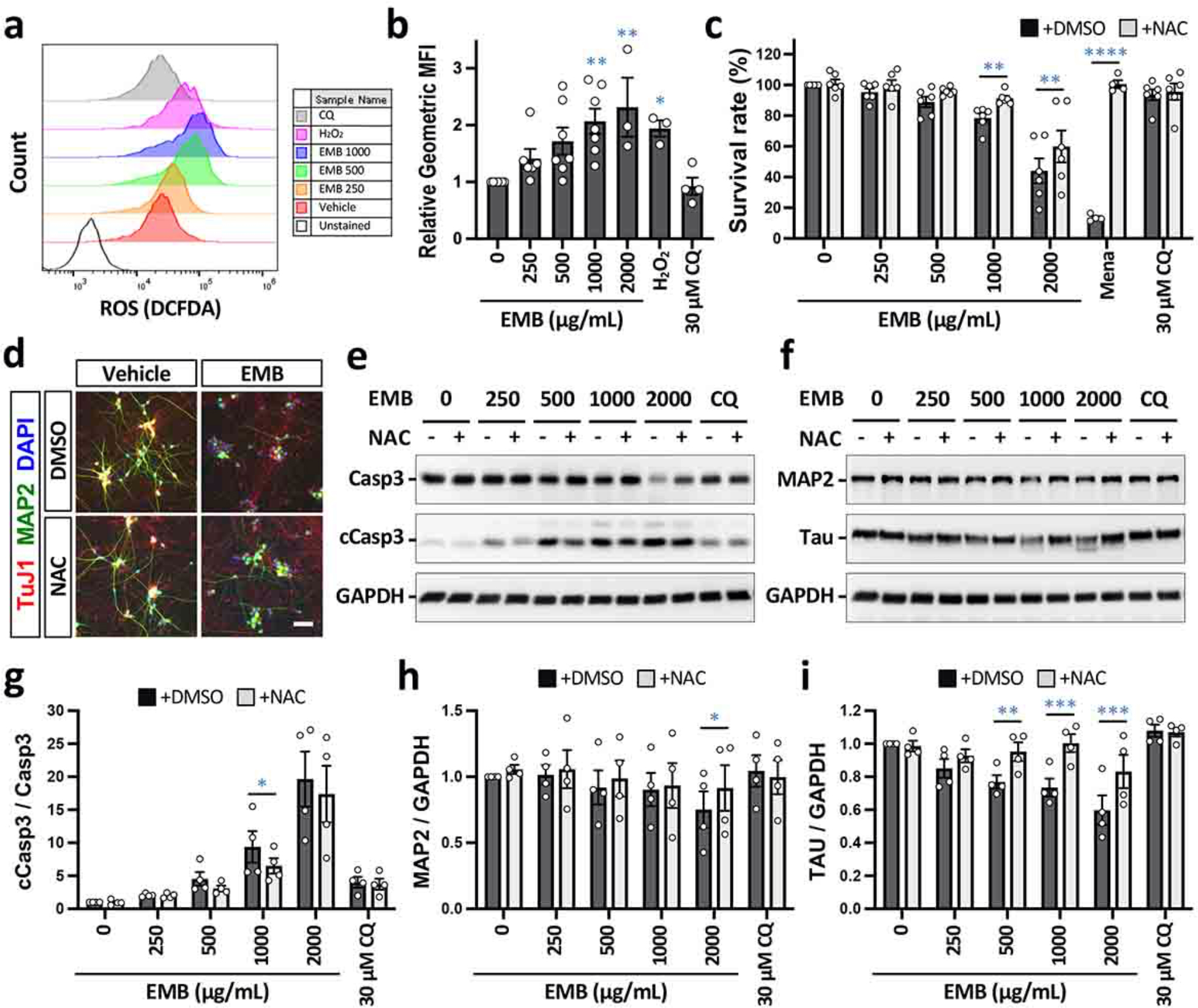
EMB-induced apoptosis and neurite degeneration are associated with reactive oxygen species (ROS) generation in iRGC-*ATOH7*/*BRN3B/SOX4* cells. (**a**) iRGC cells were treated with various concentrations of EMB for 24 hrs. The ROS generation that occurred in EMB-treated cells was measured by flow cytometry using DCFDA staining. Treatment with H_2_O_2_ for 1 hr was used as a positive control. (**b**) Quantitative results of the geometric mean fluorescence intensity (MFI) of treated samples relative to control samples and non-treated cells. All results are presented as the mean ± SEM (n = 4-7 independent experiments; *: *p* < 0.05, **: *p* < 0.01 by one-way ANOVA and Dunnett’s test). (**c**) iRGC cells were retreated with 500 μM NAC for 1 hr and treated with various concentrations of EMB for 24 hrs. Cell viability assessment was carried out using the alamarBlue® assay and is shown in the histogram. Values are presented as mean ± SEMs from four independent experiments (**: *p* < 0.01, ***: *p* < 0.001, ****: *p* < 0.0001 versus control by two-way ANOVA and Šídák’s multiple comparisons test). (**d**) The morphological changes in the iRGCs after 1000 μg/mL EMB treatment in the presence or absence of 500 mM NAC for 24 hrs and immunolabeled for MAP2 and TuJ1. Scale bar, 50 μm. (**e**) Expression levels of the apoptotic marker, caspase-3, in iRGCs at D21 using Western blotting. Glyceraldehyde 3-phosphate dehydrogenase (GAPDH) was used as a protein loading control. (**f**) Protein expression levels of MAP2 and Tau in iRGCs at D21 using Western blotting. (**g**) Quantification of the protein levels in (**e**). The intensity of each signal was normalized against the control (EMB = 0; NAC = 0). Data represent mean ± SEM (n = 4 independent batches of differentiation; *: *p* < 0.05 by two-way ANOVA and Šídák’s test). (**h** and **i**) Quantitative of the protein levels in (**f**). The intensity of each signal was normalized against the control (EMB = 0; NAC = 0). Data represent mean ± SEM (n = 4 independent batches of differentiation; *: *p* < 0.05, **: *p* < 0.01, ***: *p* < 0.001 by two-way ANOVA and Šídák’s test).

To gain further insight into the mechanism by which NAC attenuates the EMB-induced cell death in iRGC cells, we examined the levels of two autophagic markers, p62 and LC3 by Western blot analysis. As shown in Fig. 7, NAC notably suppressed the EMB-induced elevation of p62. Likewise, the levels of LC3-II were significantly reduced in the EMB/NAC co-treated group compared with the EMB-treated group. However, in a high-dose EMB treatment (2000 μg/mL for 24 hr), NAC alone was not potent enough to reduce the elevation in p62 and cleaved caspase-3 levels, indicating that the damage caused by EMB at higher concentrations in the iRGCs cannot be reversed by simply reducing the increase in intracellular ROS. It is also worth noting that NAC was not able to diminish CQ-induced autophagy deficiency, suggesting that EMB-induced generation of ROS, in turn, partially leads to the autophagy impairment in the iRGCs via a pathway that is different to that of CQ. Although it is highly likely that factors other than ROS contribute to the EMB-induced autophagy dysregulation, our results highlight a promising direction for the treatment of EMB ocular toxicity, namely one that use a combination of ROS scavengers acting as pharmacological inhibitors of ROS generation. These may be useful as a novel therapeutic remedy for EMB-induced neuropathy of RGCs when treating tuberculosis (Fig. 7d).

**Fig 7.**
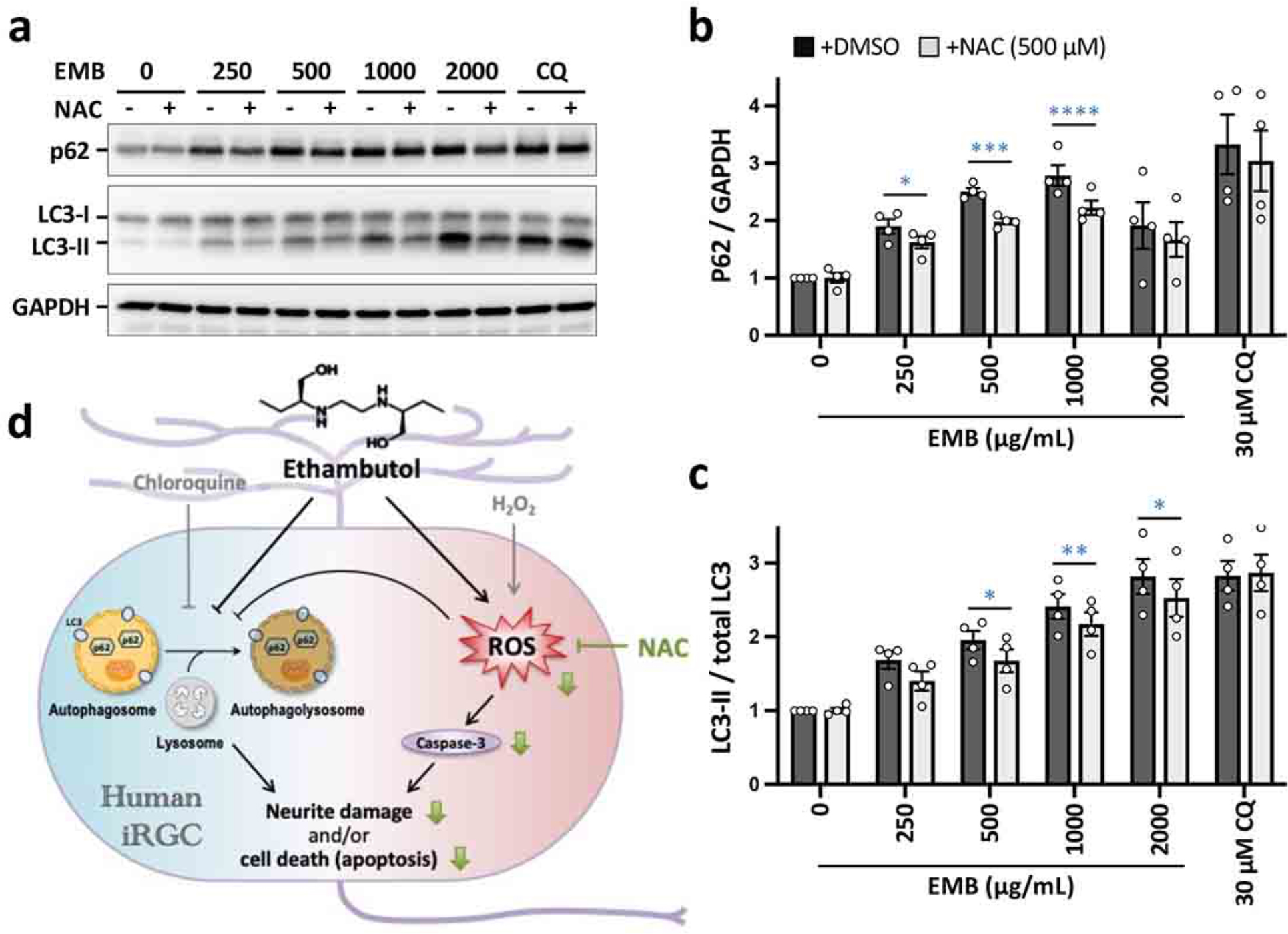
EMB-induced impaired autophagic flux was partially reversed by a ROS scavenger in iRGC-*ATOH7*/*BRN3B/SOX4*. (**a**) iRGC cells were pretreated with 500 μM NAC for 1 hr and then treated with various concentrations of EMB for 24 hrs. The conversion of LC3-I to LC3-II and p62 accumulation after EMB treatment for 24 hrs were analyzed by Western blotting. Blots are representative of the four independent experiments and GAPDH was used as th sample loading control. (**b** and **c**) Quantitative analysis of p62 (**b**) and LC3-II (**c**) levels relative to GAPDH and total LC3, respectively, were performed and the values were normalized against the control group (EMB = 0; NAC = 0). Data represent mean ± SEM of readings from 4 independent experiments (*: *p* < 0.05, **: *p* < 0.01, ***: *p* < 0.001, ****: *p* < 0.001 by two-way ANOVA and Šídák’s test). (**d**) Schematic representation of the proposed mechanisms of EMB-induced neurite degeneration and apoptosis in human RGC cells. In this study, we demonstrated that EMB inhibited autophagy flux and induced ROS generation, which then causes neurite degeneration and caspase-dependent apoptosis. Importantly, the ROS scavenger NAC blocked EMB-induced ROS generation and markedly increased neurite outgrowth and cell survival. Altogether, a combination treatment of EMB with ROS scavengers, pharmacological inhibitors of ROS generation, may be a novel therapeutic approach for the treatment of EMB-induced neuropathy in RGCs.

## DISCUSSION

Human iPSCs represent a non-ethically disputed and renewable cellular source of cells that can be used for RGC generation; this creates a unique opportunity to develop *in vitro* RGC models for various RGC-related diseases, as well as developing their use for stem cell-based regenerative medicine. In this study, we have established an efficient method of generating human RGCs from hiPSCs by the forced expression of a combination of reprogramming factors, *ATOH7*, *BRN3B* and *SOX4*. The reprogrammed iRGCs show robust expression of various RGC-specific markers, such as BRN3A, EBF1, ISL1 and RBPMS (Figs. 1, 2 and 3). We also carried out a functional assessment and demonstrated that these iRGCs display stimulus-induced channel activities and related electrical properties. The scale of production and the high yield obtained suggest this iRGC model is suitable for investigation optic neuropathies, as well as helping in the search for potential drugs for the treatment and/or prevention of optic neuropathies.

Many successes in neuronal reprogramming have been achieved using iPSCs, and RGCs are no exception. Apart from the conventional strategy of inducing RGC differentiation using small molecules and growth factors, TFs have also been utilized to accelerate the process. Overexpression of *Pax6* has been shown in mouse iPSCs to push undifferentiated stem cells first into retinal progenitor cells (RPCs), and then by subsequent differentiation into a mixed population of putative RGCs and photoreceptors [32]. A more recent report has demonstrated the use of DKK1, noggin, and Lefty A to first, induce hiPSCs into RPCs, which was followed by overexpression of *ATOH7* to further promote differentiation into RGCs [7]. While previous studies have focused on the combination of small molecules and TFs to direct a step-wise retinal neuronal differentiation, we outline in this study a faster and simpler method of achieving cell differentiation of hiPSC cells into RGCs by the direct expression of defined TFs.

Among all of the tested TFs during the search for the most efficient combination of TFs to use for RGC generation, the combination *ATOH7/BRN3B/SOX4* was found to be the most successful when driving RGC differentiation. Our iRGCs generated by this method resulted in the expression of many commonly recognized markers of RGCs, as well as the induction of various basic neuronal functions, such as stimulus-induced channel activities, as well as spontaneous electrophysiological properties. There remains a debate over whether the expression of a few RGC markers is a reliable validation of RGC identity, and therefore having certainty on the characteristics of our iRGCs remains a challenge; this is because, apart from *OPN4*, which is a specific marker of intrinsically photosensitive RGCs, there are no other markers that specifically label a cell as a RGC. In the future, the transplantation and incorporation of these iRGCs into visual circuits in an *in vivo* setting ought to further validate the identity and functionality of the cells.

Our iRGC model has proved to be suitable as a platform for the investigation of drug-induced optic neuropathy. EMB, a first-line drug used for the treatment of TB, is a common cause of toxic optic neuropathy. Here we confirmed the adverse effects of EMB on human iRGCs. EMB had a dose-dependent effect that impaired autophagic flux in iRGC-*ATOH7/BRN3B/SOX4* and it did this by blocking lysosome-autophagosome fusion, which is consistent with the results from previous studies using a mouse photoreceptor cell line RGC-5 [14]. Furthermore, we found that EMB treatment also significantly enhances ROS levels in a concentration-dependent manner. EMB is known to be a metal chelator that targets intracellular zinc, and a decrease in metal levels may have a disruptive effect on oxidative phosphorylation and mitochondrial functionality by interfering with iron containing complex I and copper containing complex IV [9, 36]. Although further confirmation is needed, interference in mitochondrion functions by EMB is highly likely to be the underlying reason for the generation of ROS and a cascade of sequential events that result in autophagy dysregulation and ultimately in apoptosis of RGCs. It is also worth noting that as EMB is known to exert a similar effect on the RGC-5 cell line, it would seem likely that the drug-induced neuropathic effect of EMB is not RGC-specific. Various other types of neurons within the neural retina, such as amacrine cells and photoreceptors, might also be susceptible to the drug’s toxic effects.

By creating our differentiation method, human iRGCs can now be generated rapidly and are suitable for drug screening test and mechanistic studies. Furthermore, the development of a range of iPSC-based technologies may help the development of novel patient-specific gene based techniques for RGC-related diseases that will help our exploration of specific pathogenetic mechanisms, as well as identifying new therapeutic interventions involving gene/cell therapy.

### Conclusions

Our hiPSC-iRGC differentiation strategy provides a simplified and efficient procedure for generating functional iRGCs in a relatively short period of time. These iRGCs will be able to serve as a human RGC model for the investigation of RGC-related diseases, as well as providing the foundation of future RGC treatments and RGC regeneration.

## DECLARATIONS

### Ethics approval and consent to participate

The uses of human iPSC lines followed the Policy Instructions of the Ethics of Human Embryo and Embryonic Stem Cell Research guidelines in Taiwan. In addition, approval from the Institutional Review Boards of National Yang Ming Chiao Tung University was obtained (#YM110195W).

### Consent for publication

Not applicable

### Availability of data and materials

All data generated or analyzed during this study are included in this article and its supplemental files. The RNA-seq data will be deposited in the Gene Expression Omnibus database. Requests for material should be made to the corresponding authors.

### Authors’ contributions

R. H.-C. Liou, S.-W. Chen and Y.-H. Wong designed and conducted most of experiments and performed the data analyses. P.-C. Wu performed the Ca^2+^ imaging experiments. H.-C. Cheng, Y.-F. Chang, A.-G Wang and M.-J. Fann initiated the hiPSC-RGC project and provided materials. R. H.-C. Liou, S.-W. Chen and Y.-H. Wong wrote the main manuscript. All the authors discussed the results, gave feedback and approved the final manuscript.

## Acknowledgements

We would like to acknowledge the grant supports from NSTC and MOE in Taiwan for this research project. We thank the Human Disease iPSC Service Consortium (funded by the Ministry of Science and Technology (MOST), MOST 108-2319-B-001-004) for iPSC generation and technical support. We also thank the Clinical and Industrial Genomic Application Development Service Center of National Core Facility for Biopharmaceuticals, Taiwan (MOST 108-2319-B-010-001) for the Sanger sequencing.

## LIST of ABBREVIATIONS

ADOA: autosomal dominant optic atrophy
BDNF: brain-derived neurotrophic factor
bFGF: basic fibroblast growth factor
CNTF: ciliary neurotrophic factor
CQ: chloroquine
DEG: differentially expressed gene
EB: embryoid body
EMB: ethambutol
hiPSC: human induced pluripotent stem cell
LHON: Leber’s Hereditary Optic Neuropathy
NAC: N-acetyl-L-cysteine
NGN2: neurogenin 2
PCA: principal component analysis
RGC: retinal ganglion cell
ROS: reactive oxygen species
rtTA: reverse tetracycline-controlled transactivator
TetO: Tet operator
TF: transcription factor
TTX: tetrodotoxin

## SUPPLEMENTARY MATERIALS

### SUPPL FIGURE LEGENDS

**Fig S1. Candidate factors for RGC induction from hiPSCs.**

(**a**) Schematic representation of the strategy used to test the candidate transcription factors (TFs) for induction of RGC differentiation from hiPSCs (upper panel). Seven TFs were selected as the candidates based on their functional roles in developmental retinogenesis and the maintenance of RGCs *in vivo* (lower panel). (**b**) Overexpression of the genes of interest was confirmed measuring protein levels using Western blot analysis after doxycycline induction. (**c**) Representative phase contrast images that display the morphological changes in the hiPSCs after induction with the indicated genes on days 1, 3 and 6. Most cells failed to survive under neuronal culture conditions after days 6. Scale bar, 20 μm.

**Fig S2. Ectopic expression of candidate TFs in hiPSCs induces neuronal-like morphology.**

(**a**) Overexpression of the genes of interest was confirmed by measuring protein levels using Western blot analysis after doxycycline induction. (**b**) Ectopic expression of TFs in hiPSCs induces neuronal-like morphology. Representative phase contrast images that display the morphological changes in the hiPSCs after induction with the indicated genes on days 1, 3, 8 and 15. The cells, on 15 days after doxycycline induction, displayed distinct neurite-like structures and were immunostained with the neuronal marker SMI312. Scale bar, 20 μm.

**Fig S3. Differentiation of hiPSC-derived RGC-like cells.**

(**a**) Flow diagram of the generation of induced RGC (iRGC). (**b**) Representative images of iRGC cells immunostained for neuronal (MAP2) and RGC (ISL1, BRN3A) markers. Scale bar, 20 μm. (**c**) Quantification of the BRN3A^+^ cells present among hiPSCs ectopically expressing the candidate genes. Data are presented as means ± SEM (n = 5 batches of independent differentiation). (**d**) Quantification of the ISL1^+^ cells present among hiPSCs ectopically expressing the candidate genes. Data are presented as means ± SEM (n = 5 batches of independent differentiation). (**e**) Quantification of the BRN3A^+^ISL1^+^ cells present among hiPSCs ectopically expressing the candidate genes. Data are presented as means ± SEM (n = 5 batches of independent differentiation).

**Fig S4. Comparison of different protocols for generation of hPSC-derived RGCs and human fetal retina.**

(**a**) A comparison of different protocols used for generating hPSC-derived RGC-like cells is presented. In our protocol, stable hiPSC lines carrying both pTetO-ATOH7-T2A-BRN3B-T2A-Puro and pTetO-SOX4-Zeo were induced for RGC-like cell differentiation upon treatment with doxycycline. Total RNA was extracted from hiPSC-derived RGCs at indicated time points and subjected to total RNA sequencing (RNA-seq). In the protocols of Gill et al. (2016) and VanderWall et al. (2020), hPSCs or hESCs were first differentiated into neuralized embryonic bodies (EB), followed by a switch to RGC maturation media containing N2 and B27 supplement. After sorting via magnetic anti-THY1 microbeads, these cells were kept in culture for 10 or 15 days before undergoing RNA-seq analysis. (**b**) The 9 datasets used for bioinformatic analysis are described.

**Fig S5. Marker gene expression and enrichment analysis in all samples.**

(**a**) A heatmap showing the expression of marker genes during retinal development and within retinal cell types in all samples. The Tet-on inducible genes are marked with asterisk (*). (**b**) A gene enrichment analysis was performed on the top 500 differentially upregulated genes (sorted by adjusted p-value and a fold change of ≥2). The eleven most significant groups are presented in terms of GOBP (left panel) and the tissue protein expression (right panel). Gray bars indicate no significant expression (padj > 0.05).

**Fig S6. EMB-induced apoptosis and neurite degeneration in iRGC-*ATOH7*/*BRN3B/SOX4* cells is able to be rescued by NAC.**

(**a**) iRGC cells were treated with various concentrations of EMB in the presence or absence of 500 μM NAC for 24 hrs. ROS generation was measured by flow cytometry using DCFDA staining. (**b**) Quantitative results of the geometric mean fluorescence intensity (MFI) of the treated cells relative to control non-treated cells. All results are presented as the mean ± SEM (n = 4 independent experiments; *: *p* < 0.05 by two-way ANOVA and Šídák’s test). (**c**) The morphological changes in the iRGCs after 1000 μg/mL EMB treatment in the presence or absence of 500 μM NAC for 24 hrs and immunolabeled for MAP2 and TuJ1. Scale bar, 50 μm.

